# Using long-read sequencing to detect imprinted DNA methylation

**DOI:** 10.1101/445924

**Authors:** Scott Gigante, Quentin Gouil, Alexis Lucattini, Andrew Keniry, Tamara Beck, Matthew Tinning, Lavinia Gordon, Chris Woodruff, Terence P. Speed, Marnie E. Blewitt, Matthew E. Ritchie

**Affiliations:** The Walter and Eliza Hall Institute of Medical Research, 1G Royal Parade, Parkville, 3052 Australia; Department of Medical Biology, The University of Melbourne, Parkville, 3010 Australia; Australian Genome Research Facility, 305 Grattan Street, Melbourne, 3000 Australia; School of Mathematics and Statistics, The University of Melbourne, Parkville, 3010 Australia; Department of Genetics, Yale University, New Haven, 06520 USA

**Keywords:** Nanopore sequencing, Differential methylation, Haplotyping, Imprinting, Long-read sequencing

## Abstract

Systematic variation in the methylation of cytosines at CpG sites plays a critical role in early development of humans and other mammals. Of particular interest are regions of differential methylation between parental alleles, as these often dictate monoallelic gene expression, resulting in parent of origin specific control of the embryonic transcriptome and subsequent development, in a phenomenon known as genomic imprinting. Using long-read nanopore sequencing we show that, with an average genomic coverage of approximately ten, it is possible to determine both the level of methylation of CpG sites and the haplotype from which each read arises. The long-read property is exploited to characterise, using novel methods, both methylation and haplotype for reads that have reduced basecalling precision compared to Sanger sequencing. We validate the analysis both through comparison of nanopore-derived methylation patterns with those from Reduced Representation Bisulfite Sequencing data and through comparison with previously reported data.

Our analysis successfully identifies known imprinting control regions as well as some novel differentially methylated regions which, due to their proximity to hitherto unknown monoallelically expressed genes, may represent new imprinting control regions.

## Introduction

Methylation of the 5th carbon of cytosines (5mC or simply mC) is an epigenetic modification essential for normal mammalian development. Methylation differences between alleles contribute to establishing allele-specific expression patterns. As obtaining genome-wide haplotyped methylomes with short reads remains challenging, we evaluated the ability of long read, nanopore-based sequencing to improve allele-specific methylation analyses.

We apply the technique to the study of genomic imprinting, where differential expression of the maternal and paternal alleles in the offspring is at least partially set by the differential methylation (1–5). Imprinting is proposed to arise from the diverging interests of the maternal and paternal genes (6). In accordance with its primordial role in allocation of resources from the mother to the offspring, the placenta, along with the brain, is the organ where parental conflict results in the most pronounced imprinted expression (7–9). We thus conduct a survey of differential methylation and expression in murine embryonic placenta.

Recent studies have increased the number of genes identified as subject to imprinting in mouse to about 200 (10–15). The cause of the differential expression between paternal and maternal alleles is only known for a subset of these genes; maternal histone marks can play a role (14), and in other cases it involves the differential methylation of adjacent regions (5). The differential methylation patterns may be established in the gametes and persist through the epigenetic reprogramming occurring after fertilisation (16). These differentially methylated regions (DMRs) are called primary DMRs, or imprinting control regions (ICRs). Alternatively, differential methylation may arise during development, perhaps as a downstream effect of differential expression, in which case the regions are called somatic or secondary DMRs (17).

Apart from the parent of origin of the allele, genetic differences can also be associated with differential methylation. In this case, F1 hybrids of distinct mouse strains will display DMRs between the alleles according to the strain of origin (18), and not the parent. Genetically determined DMRs can have profound effects on phenotype, for instance in humans by altering the expression of mismatch repair genes important in cancer (19). Therefore, we also investigate the link between DNA methylation and expression for strain-biased genes.

Reconstructing haplotyped methylomes necessitates the simultaneous measurement of DNA methylation and single-nucleotide polymorphisms (SNPs) differentiating the alleles. This can be achieved by deep sequencing of bisulfite-converted DNA on the Illumina platforms, although the short reads combined with the reduced complexity of the bisulfite-treated DNA make the process inefficient, meaning many regions with low SNP density remain unresolved. Long reads provided by third generation sequencing technologies can overcome the requirement of a high SNP density, while several methods allow the assessment of base modifications on native DNA (thus also avoiding the reduction in complexity associated with bisulfite conversion). These methods include: analysis of polymerase kinetics for PacBio SMRT sequencing (20), and detection of deviations in the electric signal for Oxford Nanopore sequencing, via nanopolish (21), signalAlign (22), mCaller (bioRxiv doi:10.1101/127100), Tombo (bioRxiv doi:10.1101/094672), or DeepSignal (bioRxiv doi:10.1101/385849). We note that, for the dominant eukaryotic genome base modification at 5mC, the PacBio technology requires very high coverage making it impractical for use in the analysis of mammalian genomes (23). PacBio SMRT sequencing can be combined with bisulfite treatment (SMRT-BS) to facilitate 5mC detection, but this approach is currently only available for targeted sequencing (24) and the bisulfite treatment introduces the same drawbacks noted above in addition to fragmenting the DNA. Additionally, while PacBio technology is limited to maximum read lengths of between 50 and 100 kb (25), Oxford Nanopore sequencing has no theoretical upper limit on read length and exhibits no bias in sequencing quality with read length (26), which is especially beneficial i n genomic regions devoid of SNPs, or highly repetitive regions.

Here we use the MinION and PromethION long-read nanopore sequencers to generate whole-genome haplotyped methylomes from murine embryonic placenta. With a mean coverage of 10X we successfully identify known imprinting control regions as well as novel parent-of-origin DMRs near imprinted genes, as well as strain-specific DMRs close to both strain-biased genes and structural variants. We show the improved efficiency of this strategy over existing workflows to resolve allele-specific methylation, and highlight its utility in investigating the mechanisms of genomic imprinting.

## Results

### Nanopore methylation calls are concordant with other technologies

We sequenced the embryonic portion of placenta derived from a male embryonic day 14.5 (E14.5) conceptus from a C57BL/6 (Black6, or B6)× *Castaneus* (Cast) F1 on the MinION platform to a depth of about 8X (Fig. S1) and called methylation using *nanopolish* (21). The genome-wide methylation data successfully recapitulated known patterns: CpG islands (CGIs), as defined by Irizarry *et al.* (27), separated into two groups of high and low methylation (Fig. 1A); the methylation level dipped at transcriptional start sites (TSSs) (Fig. 1B), and the average genome-wide methylation level was around 50%, as previously reported for placental tissue (28).

**Fig. 1.**
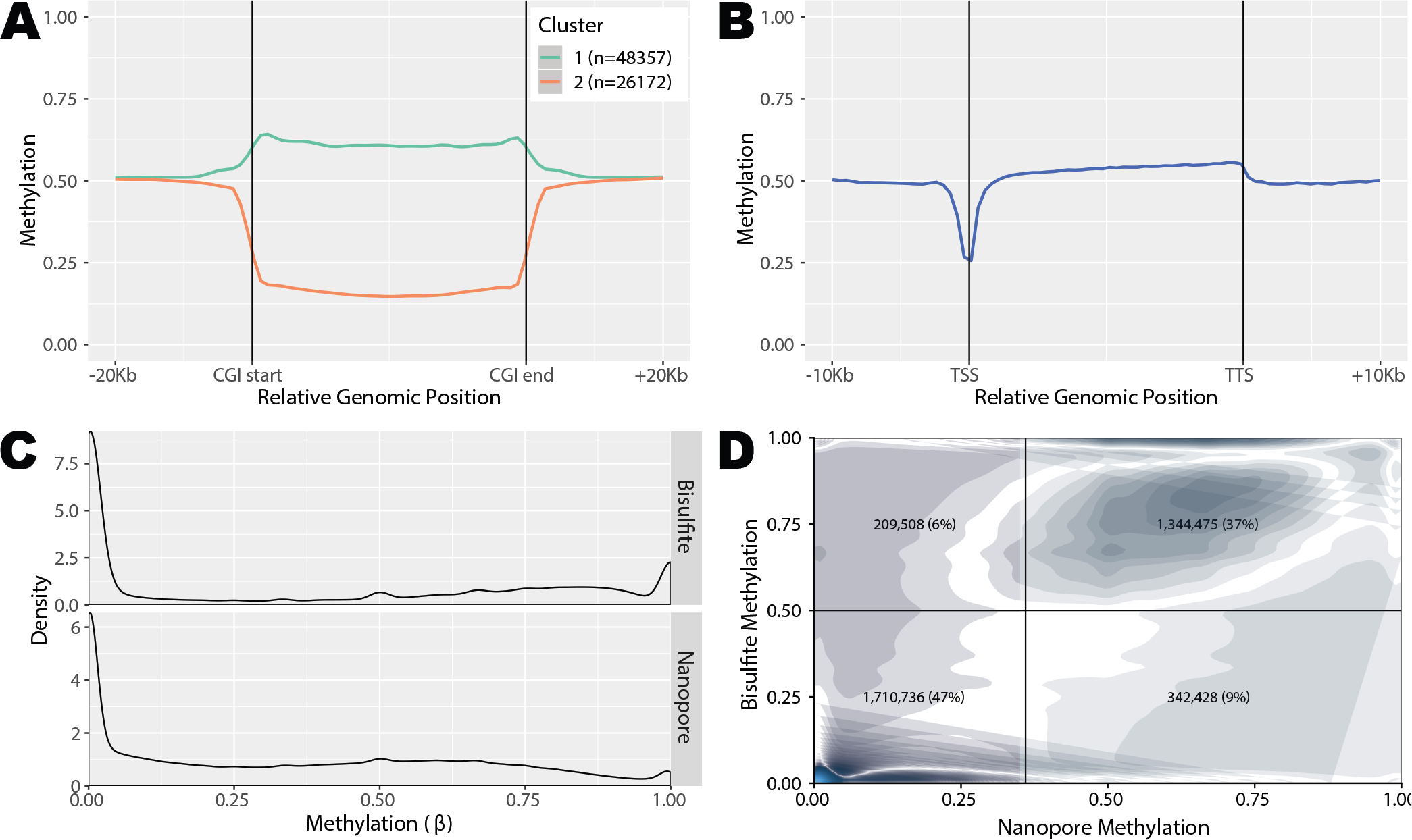
Nanopore methylation calls are consistent with expected results and established technologies. **A.** Metaplot of nanopore methylation calls across CpG islands, clustered in two groups of high and low methylation. **B.** Metaplot of nanopore methylation calls across the aggregated gene bodies of all protein-coding genes recapitulated known methylation structures. **C.** Density of methylation calls (*β*, the average methylation based on all reads covering that position) for sites covered by both nanopore and RRBS. **D.** Joint density of nanopore and RRBS methylation calls for the same sites as in C. Darker regions indicate regions of higher density, while lighter regions indicate regions of lower density. The density plot is split into four quadrants according to a RRBS threshold of 0.5 and a nanopore threshold of 0.36, and the percentage of sites in each quadrant is displayed.

To further validate the accuracy of the nanopore methylation calls, we compared them to Reduced Representation Bisulfite Sequencing (RRBS) data on the same sample at sites covered by both methods. Nanopore methylation calls showed an overall similar distribution to RRBS methylation calls, albeit with a bias toward intermediate values of methylation (Fig. 1C). Despite being less correlated than measurements from methylation-specific technologies (29) with a median absolute deviation (MAD) of 0.18, per-site methylation was also relatively concordant between the two methods, with 85% of CpG sites being called correctly when converting average methylation values to binary calls. (Fig. 1D).

Sequencing of E14.5 embryonic placenta from the reciprocal cross (Cast × B6) on the PromethION platform at 12X coverage (Fig. S1) generated comparable results.

### Increased haplotyping efficiency with nanopore reads

We next used high-confidence SNPs between the Cast and B6 strains to haplotype RRBS and nanopore reads. In order to mitigate the high sequencing error rates of nanopore sequencing, we used two methods of haplotype assignment, denoted the ‘*basecall method*’, based on FASTQ data, and the ‘*signal method*’, based on the *phase-reads* module from *nanopolish*, an HMM which uses the raw nanopore signal to predict genotype (30). Additionally, we only assigned a haplotype to those nanopore reads with at least five high-confidence SNPs (Fig. S3). All RRBS reads overlapping at least one SNP were assigned a haplotype (31). Only 24% of the mapped RRBS reads could be assigned to one haplotype, whereas 74% of mapped nanopore reads were haplotyped in the expected proportions (Fig. 2A and B): roughly half of the haplotyped reads were assigned to the maternal haplotype, and half to the paternal haplotype, albeit showing a slight bias towards the paternal haplotype (due to an increased number of split reads in regions where sections of the Cast genome has a deletion with respect to the B6 genome). The pattern of haplotype assignment was consistent across the autosomal chromosomes, while, as expected for a male sample, almost all (91%) of the reads aligned to the X chromosome were assigned to the maternal haplotype (Fig. 2B). Haplotyping of the Cast × B6 cross gave similar results (Fig. S2A). The lack of maternal bias in read haplotype indicates minimal maternal contamination, which is also reflected in consistent RNA-seq library sizes (Fig. S4).

**Fig. 2.**
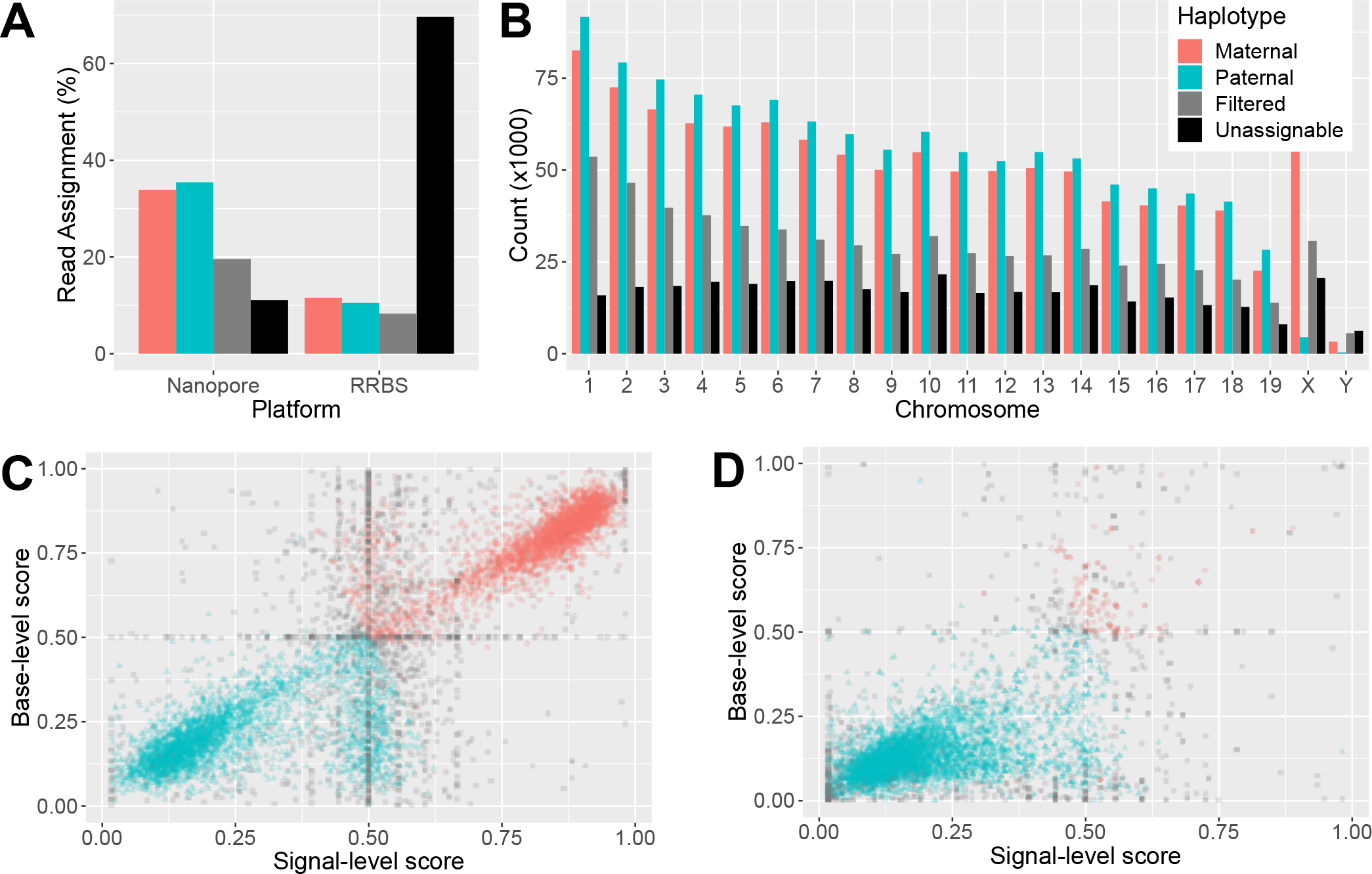
Accurate and efficient haplotyping of nanopore reads. **A.** Percentages of mapped reads from RRBS and nanopore sequencing that were assigned to the B6 genome (maternal), Cast genome (paternal), or that could not be haplotyped (filtered) for the B6 × Cast F1 sample. **B.** Percentages of mapped reads from nanopore sequencing that were assigned to each haplotype on each chromosome. **C.** Scatter plot of haplotype scores for nanopore reads according to signal (x-axis) and basecall (y-axis) methods. Only 10,000 randomly selected reads are shown for ease of visualisation. **D.** Signal and basecall haplotype scores for reads from the sequencing of the pure parental Cast strain.

We further evaluated the accuracy of the haplotyping of the nanopore reads by sequencing the same tissue from the parents (B6 only, and Cast only). Following the same haplotyping procedure, 85.7% of the reads were correctly assigned to the relevant genotype, and 1.5% were misassigned (Fig. S2B and C).

The large majority of nanopore reads showed strong agreement between the two haplotyping methods. Discrepancy between the basecall and signal methods are typically due to a low number of SNPs being scored by one or both methods, resulting in these alignments being filtered out by the haplotyping procedure (Fig. S3). However, when examining the overall predictive performance of the two methods with AUROC (Area Under Receiver Operating Characteristic Curve) on the single-strain experiments, the signal method marginally outperformed the basecall method (see Table 1). We also found that the combination of the two methods achieved a slight improvement again over the signal method, either with a logistic regression model (32) or an *ad hoc* combination of the two approaches (see Methods). The *ad hoc* method allowed classification of an 30,000 additional reads over the logistic regression and signal method, both of which excluded reads for which *nanopolish* failed to produce output.

**Table 1.** Accuracy and support of haplotyping methods on pure-strain reads.

| Method | AUROC | Accuracy (%) | Called Reads |
| --- | --- | --- | --- |
| Basecall | 0.976 | 0.963 | **261,806** |
| Signal | 0.988 | 0.961 | 199,474 |
| Regression | **0.992** | 0.972 | 198,684 |
| *Ad hoc* | — | **0.983** | 228,866 |

In both nanopore and RRBS data, the main cause of haplotyping failure is the lack of SNPs in the region covered by the read (Fig. S3). While the proportion of successfully haplotyped nanopore reads could increase with optimisation for longer reads and anticipated improved sequencing accuracy, RRBS haplotyping efficiency is limited by the short read length.

### Parent-of-origin and strain-biased gene expression

To investigate the correlation between differential methylation and differential gene expression, we performed RNA-seq on the same F1 placental tissue from reciprocal crosses of B6 and Cast, in quadruplicates. Maternal tissue contamination was unlikely as for each embryo maternal and paternal counts were similar (Fig. S4). We found 135 genes with a parental bias in expression (imprinted genes, 10% FDR, Fig. 3A): 88 with higher expression from the maternal allele and 47 with higher expression of the paternal allele. Among the 135 genes, 53 corresponded to well-characterised imprinted genes in classic databases (33–36). A further 17 of these genes, including *Fkbp6*, *Smoc1/2*, *Gzmc/d/e/f/g*, *Zdhhc14* and *Arid1b* have been identified as imprinted in one or several recent studies (12–15). The remaining 65 genes constitute novel candidate genes with parent-biased expression in mouse placenta. The complete annotated list is reported in Additional File 1.

**Fig. 3.**
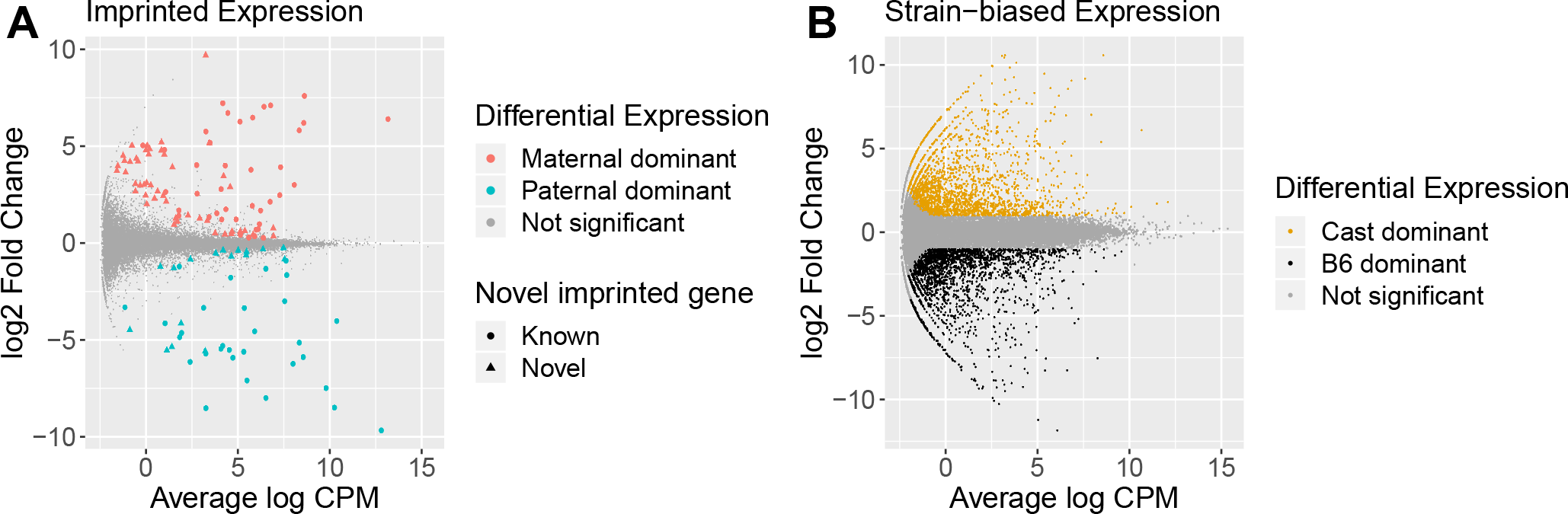
Differential allelic expression in mouse E14.5 embryonic placenta. **A.** Differential expression between the maternal and paternal alleles. Genes with adjusted p-value < 0.1 are coloured in red when maternal expression dominates (positive log-fold change) and blue when paternal expression is greater (negative log-fold change). The shape of the point indicates whether the differentially expressed gene has previously been reported as imprinted. **B.** Differential expression between B6 and Cast alleles. Genes with adjusted p-value < 0.05 and log2 fold-change > 1 are coloured in black when B6 expression is higher and orange when Cast expression is higher. Interactive plots are available at bioinf.wehi.edu.au/haplotyped_methylome.

We also identified 4,029 genes (13% of expressed autosomal genes) with a strain bias greater than two-fold (5% FDR, Fig. 3B and Additional File 1), evenly split between B6 dominance (2,027 genes) and Cast dominance (2,002).

### Known imprinting control regions are observable by nanopore sequencing

We combined the methylation and haplotyping data from the nanopore reads to compare methylation between the parental alleles. To highlight the linkage information of the methylation data available for nanopore reads, as well as the per-site per-read data, we plotted the loess fit of the cytosine methylation levels for each read in the region of interest (Fig. 4A and B).

**Fig. 4.**
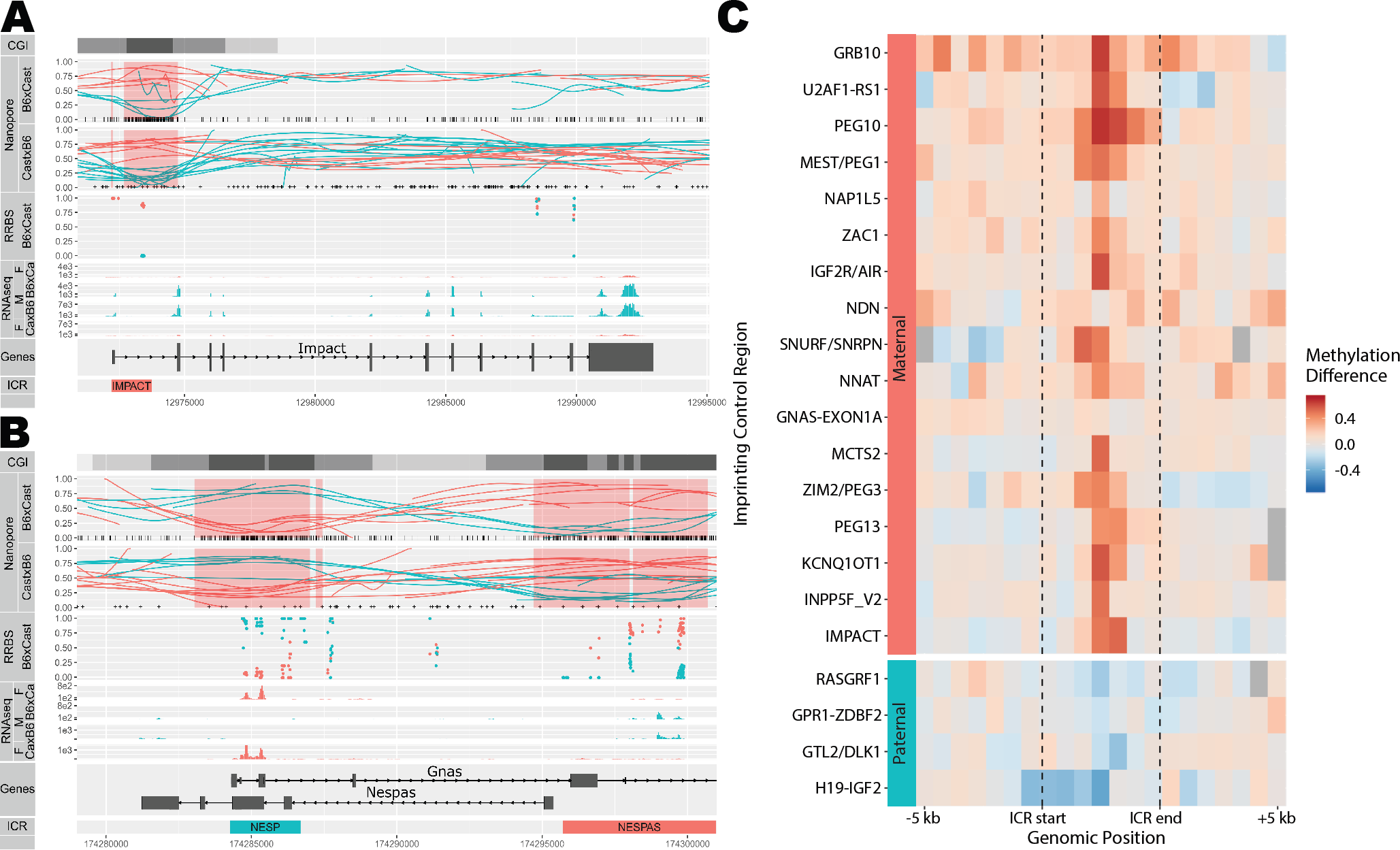
Nanopore allele-specific methylation captures known differential methylation at imprinting control regions. **A.** Allelic methylation plot of maternally imprinted gene Impact displays a clear DMR at its imprinting control region (ICR). Haplotyped RRBS data shows concordance with nanopore allelic methylation. Allele-specific RNA-seq coverage plots show monoallelic paternal expression. *CGI*: CpG Islands are displayed in black, with CpG shores in dark grey and CpG Shelves in light grey. *Nanopore*: Vertical bars at the base of the *B6Cast* track denote CpG sites used for methylation calling, while ‘+’ signs at the base of the *CastB6* track denote SNPs used for haplotyping. Highlighted red regions indicate DMRs detected by DSS. The maternal allele is shown in red and the paternal allele in cyan for all plots. **B.** Allelic methylation plot as in A. for the reciprocally imprinted genes Nespas and Gnas. RNA-seq gives very low expression and is not shown. **C.** Heatmap of differences (maternal *−* paternal) in allelic methylation in relative-width bins along known ICRs. Regions are sorted in order of average methylation difference, with regions in the same imprinting cluster placed adjacent to each other. Regions without haplotyped calls for both alleles are shown in gray.

Differentially methylated regions at known imprinting control regions (35) were readily visible and concordant with matched allele-specific RNA-seq and RRBS data (Fig. 4A and B). Nanopore data recapitulated methylation differences at most known imprinting control regions (Fig. 4C), often showing extended differential methylation past the annotated ICR borders.

### Nanopore sequencing reveals novel differentially methylated regions

Next, we sought to define DMRs between parental alleles as well as between strains *de novo*, using the differential methylation tool DSS (37). We ranked putative DMRs based on the area statistic. Using the DSS default threshold of 10^−5^, we obtained a total of 933 DMRs, of which 309 were explained by parent-of-origin differences, and the remainder by strain-specific effects (Additional File 2). We then examined these DMRs, in conjunction with haplotyped RRBS and RNA sequencing data for corroborating evidence of differential methylation and differential expression respectively, in order to find putative DMRs of interest at imprinted genes.

Of the 20 highest ranking DMRs, 15 corresponded to known ICRs. Although many of the lower-ranking DMRs are potential false-positives, they also included regions of known imprinted expression (for instance two small detected DMRs immediately adjacent to known DMRs at the IMPACT and NESP ICRs, shown in Figures 4A and B respectively.) Thus in the absence of statistically robust DMR-finding methods for nanopore data we kept this permissive threshold.

Five ICRs annotated in the WAMIDEX database (35) were not detected *de novo* (Table 2). INPP5F_V2 and GRB10 simply lacked coverage in the B6 × Cast sample but showed clear differential allelic methylation in the Cast × B6 sample; GNAS-EXON1A also lacked coverage in B6 × Cast but the reciprocal sample and the RRBS did not suggest differential methylation, while NDN lacked coverage in both samples. The last undetected region was GPR1-ZBDF2; however Duffié et al. (17) have shown that this region lacks important features of a *bona fide* ICR, and that a neighbouring maternally hypermethylated region is the true ICR. The region in question was readily detected as differentially methylated from the nanopore data (DMR #229, Table 2).

**Table 2.**
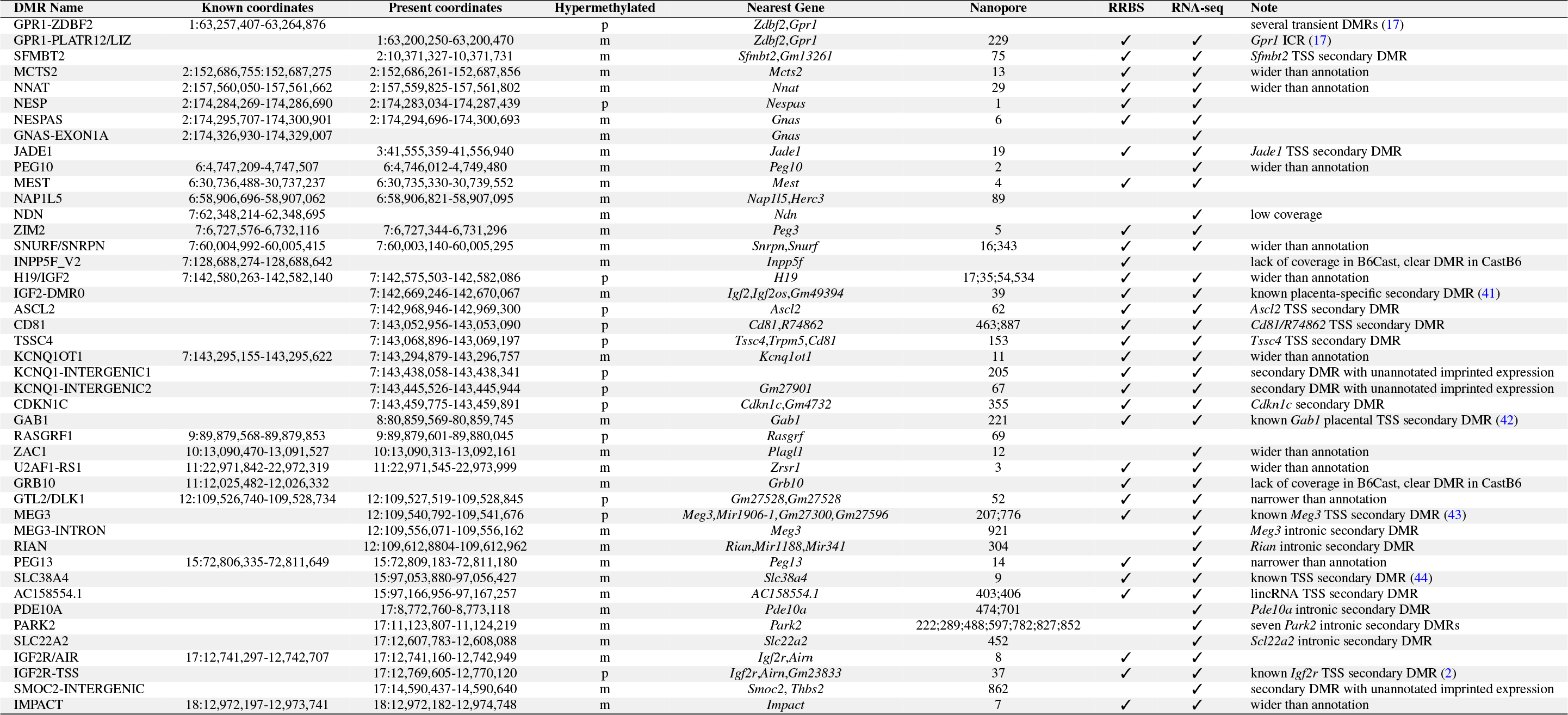
List of known and proposed DMRs associated with imprinted genes. A check mark is shown where RRBS and RNA-seq evidence supports the DMR. The DMR rank is shown when supporting nanopore data exists.

In addition to ICRs, we detected numerous DMRs at imprinted genes that do not appear to be present in gametes (38–40), and which therefore likely constitute secondary DMRs (Table 2). When RRBS coverage was present, the bisulfite data corroborated the *de novo* DMR identification. Five of the secondary DMRs have been described previously, although they are not currently compiled in a database: maternal hypermethylation at the *Igf2* promoter (41), maternal hypermethylation at the placental-specific promoter of *Gab1* (42), paternal hypermethylation of the *Meg3* transcriptional start site (TSS) (43), maternal hypermethylation at the *Slc38a4* TSS (44), and paternal hypermethylation at the *Igf2r* promoter (2).

The remaining secondary DMRs have not been previously characterised. Six of them overlapped the transcriptional start sites of imprinted genes: *Sfmbt2*, *Jade1*, *Ascl2*, *Cd81/R74862*, *Tssc4*, and *AC158554.1*.

Other novel secondary DMRs overlapped introns rather than transcriptional start sites. *Park2*, a recently identified maternally-biased gene (13), had seven intronic DMRs, all displaying hypermethylation of the maternal allele. *Rian* displayed a DMR that had not been previously reported in mice, although its human ortholog also presents an intronic imprinted DMR (45).

In some cases, inspection of the parent-specific DMRs revealed unannotated imprinted transcription nearby, for example in the *Kcnq1* and *Igf2r* clusters (Table 2). These RNA-seq reads may be part of imprinted long non-coding RNAs, frequent at imprinted clusters.

While we have collated all the imprinted DMRs that we found to directly overlap with imprinted expression -in addition to the WAMIDEX ICRs- in Table 2, we note that other imprinted DMRs may be associated with the imprinted expression of more distant genes, or with genes that are only expressed or imprinted in specific tissues. For example we found imprinted DMRs in the promoters of *Smoc2* (DMR #224) and *Arid1b* (DMR #863), two genes recently identified as being imprinted (and also imprinted in our data). The strong DMR #110 overlapped the transcriptional start site of *Gtsf2*, which was poorly expressed in placenta but may be imprinted in the tissues where it is expressed (in gonocytes and spermatids (46)). In the absence of chromatin conformation data or functional validation, we did not attempt to formally assign these DMRs to specific genes.

We however used the top, most reliable 400 DMRs to calculate the distance of genes to their nearest DMR depending on their expression status. We found that parentally biased genes were more likely to be proximal to parent-of-origin DMRs than unbiased genes (median distance 0.9 Mbp compared to 7.4 Mbp), whereas strain-biased genes and DMRs did not show this relationship (median distance 2.9 Mbp compared to Mbp) The distributions of distances to the nearest DMR are shown in Fig. 5, which shows a striking relationship between parentally biased genes and parent-of-origin DMRs. This result is consistent with parental bias in expression being driven necessarily by epigenetic differences, whereas differential expression between strains is mainly driven by genetic differences.

**Fig. 5.**
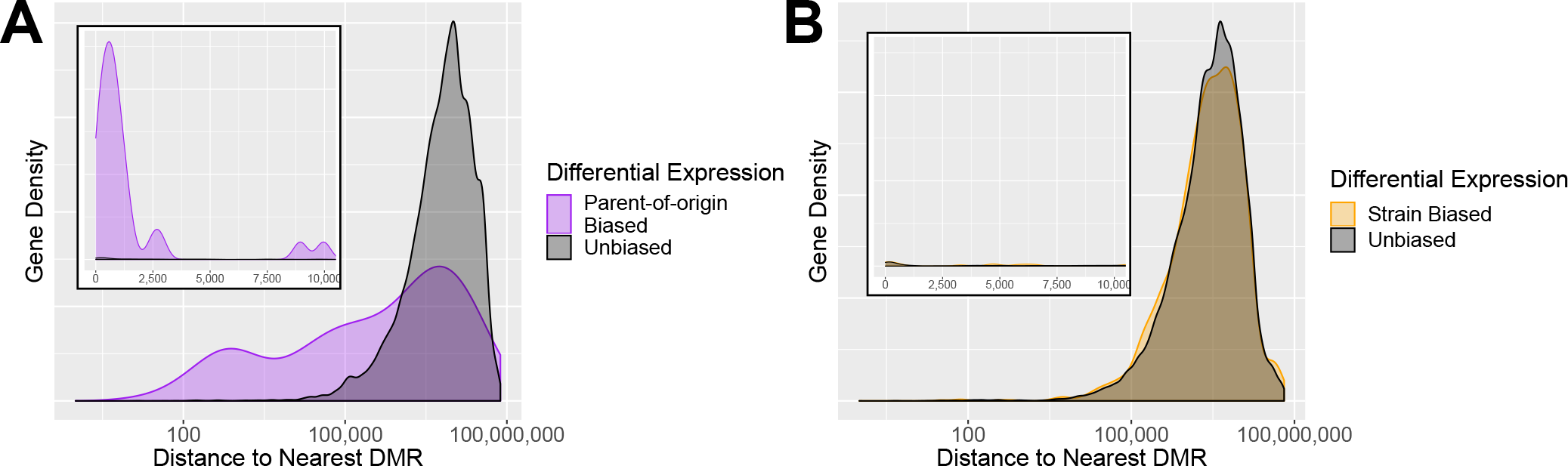
Proximity of differentially expressed genes to DMRs. **A.** Distribution of distance from genes to imprinted DMRs, shown on a log-scale. Inset shows distances from 0 to 10,000bp on a linear scale. Imprinted genes are much more frequently located within 100-100,000 bp of an imprinted DMR. **B.** Distribution of distance from genes to strain-specific DMRs. Strain-biased genes are for the most part located no closer to a strain-specific DMR than non-differentially expressed genes, indicating the strain-specific differential expression is likely caused by other factors, such as genomic differences. In both cases, we use only DMRs ranked in the top 400.

### Long reads provide advantages in differential methylation analysis

Inspection of the DMRs revealed multiple advantages of our nanopore-based method of methylome haplotyping over traditional bisulfite sequencing (Fig. 6).

**Fig. 6.**
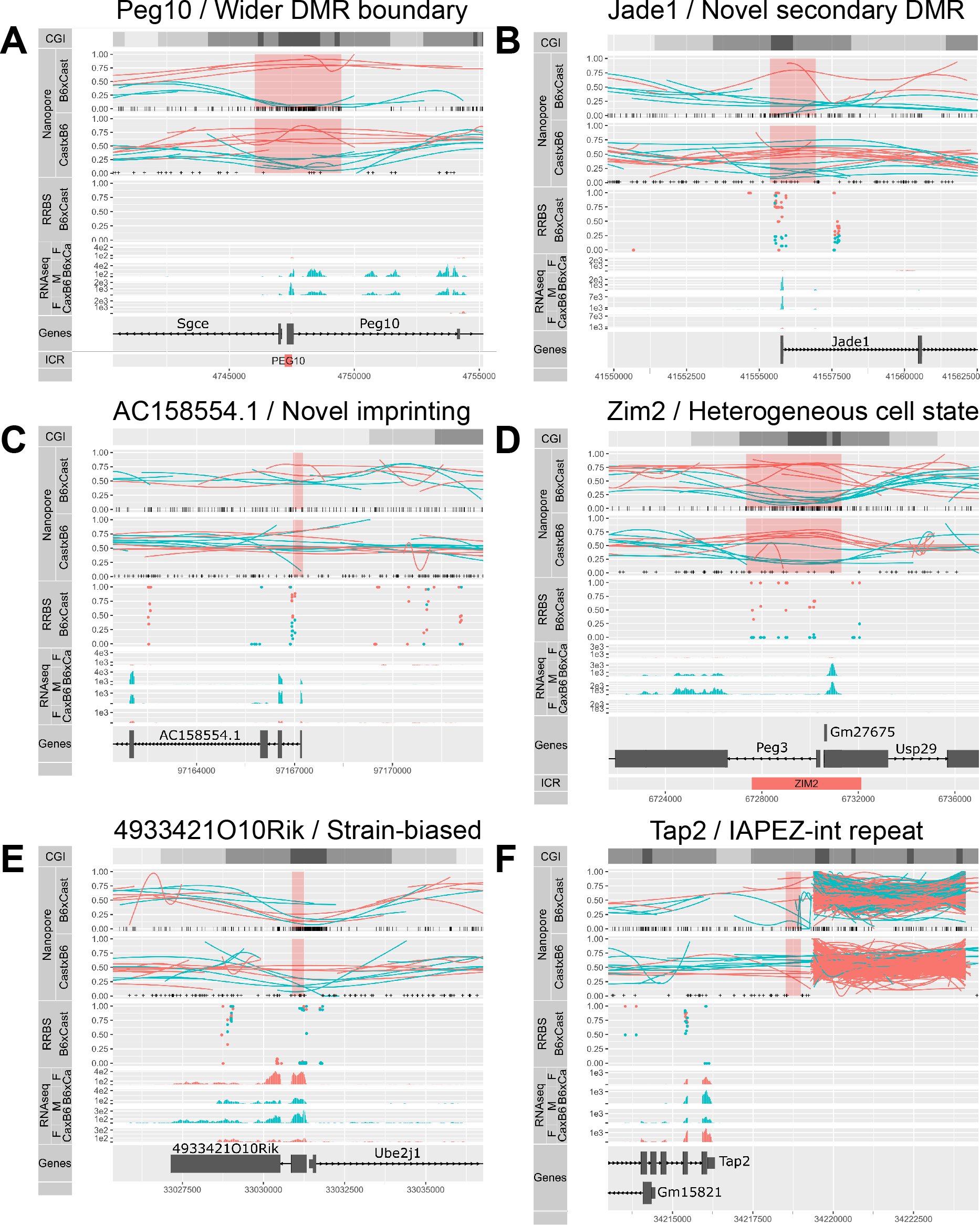
Examples of *de novo* DMRs and the advantages proffered by long reads. **A.** Allelic methylation (as in Figure 4) plot of maternally imprinted gene Peg10 displayed a clear DMR at its ICR which was much wider than the previously annotated DMR (bottom). **B.** Previously uncharacterised secondary DMR at the TSS of maternally imprinted gene Jade1. **C.** Novel maternally imprinted gene AC158554.1, with imprinted methylation at its TSS. **D.** Allelic methylation plot of maternally imprinted gene Peg3 showed consistently high methylation across some maternal reads, and consistently low methylation across others, a conclusion that could not be drawn from the middling bisulfite methylation values. **E.** Strain-of-origin DMR associated with the strain-biased expression of 493342110Rik. **F.** DMR associated with the omission of a IAPEZ repeat from the Cast genome, suggesting that the methylation in the flanking region was affected by the presence or absence of the repeat.

We were able to resolve DMRs in regions of low SNP density, where there were no haplotyped RRBS reads despite the presence of a CpG island (Fig. 6A). The particular DMR in Fig. 6A encompassed the transcriptional start site of the imprinted gene Peg10, and was much wider than the previously annotated ICR. The increased DMR width was a regular occurrence at ICRs (Table 2).

Our method also uncovered novel secondary DMRs at known imprinted genes such as *Jade1* (Fig. 6B), as well as at previously uncharacterised imprinted transcripts such as *AC158554.1*, annotated as a lincRNA (ENSMUSG00000116295, Fig. 6C).

Another advantage provided by the long reads was apparent at the ZIM2-PEG3 ICR (Fig. 6D). RRBS data from the B6Cast F1 showed that certain CG dinucleotides were highly methylated on the maternal allele (100% methylation at these positions) while others were variably methylated, resulting in averages of 25-50% methylation. Two scenarios could give rise to these intermediate values: either the variable positions are randomly unmethylated in all maternal alleles, or there exists two populations of maternal alleles, one where CG dinucleotides are methylated throughout the region and one where the variable positions are consistently unmethylated. The long nanopore reads revealed that the second scenario contributes to the observed intermediate methylation patterns: there was a mixture of cells, in some of which the maternal allele showed a contiguous loss of methylation. Although this result is well known to those who have practiced Sanger bisulfite sequencing, the haplotyped methylome derived from nanopore sequencing allowed investigation of this variability more accurately (no PCR bias) and across the whole genome.

Eight strain-specific DMRs were also found within 5 kb of a gene with strain-biased expression (Fig. 6E). Most of these exhibit structural variation proximal to the DMR, although a small number exhibited changes in expression seemingly not associated with any structural variant.

One example of a structural variant between B6 and Cast associated with differential methylation can be found on Fig. 6F. Upstream of the *Tap2* gene, an IAPEZ retrotransposon in the B6 genome is absent from the Cast genome (Cast reads are truncated upstream of the repeat and absent downstream), and the gene-proximal border is more highly methylated on B6 alleles than on Cast alleles. This differential methylation would be consistent with the insertion of the transposon attracting methylation that spreads to adjacent regions. We could also see at this repeat region a lot of spuriously mapping reads from both strains, suggesting that the repeat is present in multiple copies that the current assembly fails to account for.

## Discussion

Determining allele-specific m ethylation p atterns i n diploid or polyploid cells with short-read sequencing is hampered by the dependence on a high SNP density and the reduction in sequence complexity inherent to bisulfite treatment. In this study we demonstrate the use of long-read nanopore sequencing to derive haplotyped methylomes of the embryonic portion of mouse placentae. Methylation estimates from nanopore reads are consistent with previous knowledge (Fig. 1). The longer read lengths allowed most reads to overlap multiple SNPs, resulting in accurate haplotyping of 75% of the reads, a much higher proportion than comparable short-read data (Fig. 2). Sequencing of native DNA not only maintains the sequence complexity that is lost in bisulfite treatment, but also has the potential to detect a variety of base modifications outside 5mC, bypassing the need for specialised chemistries such as bisulfite (for 5mC) or oxidative-bisulfite (for 5 hmC) treatments. Furthermore, we are able to characterise allele-specific methylation at a relatively shallow level of genomic coverage (~10X), which is substantially lower than the coverage required by Pacific Biosciences single-molecule sequencing to ascertain any native base modification (25X) or 5mC in particular (250X) (23). Nonetheless, a detailed comparison of the performance of nanopore and PacBio in the detection of base modifications on matched samples would be of interest.

Recent increases in throughput of nanopore sequencing instruments make this approach a cost-effective way of obtaining genome-wide allele-specific methylation for mammalian-sized genomes, compared to the alternatives of short-read whole-genome bisulfite sequencing, or PacBio SMRT sequencing. Thus the approach we present is unique in its ability to characterise allele-specific single-molecule cytosine methylation state in eukaryotes, in which the 5mC modification is both common and highly relevant to transcriptional regulation.

The haplotyped methylomes for reciprocal B6 × Cast F1 samples confirm the parent-of-origin specific methylation of ICRs and provide an improved definition of their boundaries (Fig. 4). By integrating the haplotyped methylomes with allele-specific expression data, we identified novel DMRs linked to imprinted genes. These are likely to constitute secondary DMRs, whose role and origin are unclear. We note that the low sequencing coverage in this study (~ 10X) limits our ability to detect modest methylation differences between alleles, and is thus most suited to the detection of large differences such as those occurring at ICRs. We confirm a large number (70) of previously identified imprinted genes and propose another 65 as new candidates (Fig. 3 and Additional File 1). This suggests that although the monoallelically expressed genes are now well characterised, sensitive analyses can still uncover parentally biased genes. Interestingly, though we find more maternal-dominant genes than paternal-dominant ones (88 and 47, respectively), the imbalance is much less pronounced than in Finn *et al.* (2014) (12) (96% maternal dominance). Applying long-read sequencing to the transcriptome also promises improvements in the percentage of usable data, the detection of allele-specific as well as isoform-specific differential expression, and even the detection of RNA base modifications.

Our allele-specific methylation and expression data can also be used to reveal strain-biased expression of genes linked to strain-specific DMRs. The genetic divergence between the two strains accounts for most of the differences in expression, however the presence of DMRs could suggest an epigenetic component to the regulation of a subset of genes.

We foresee a number of improvements that will make the determination of haplotyped methylomes by nanopore sequencing more efficient and comprehensive in the future. Firstly, we expect to see an expansion in the types and nucleotide contexts of base modifications characterised. Our analysis is based on Simpson *et al.* (2017) (21), and is limited to 5mC at CpG sites. However that is not a limitation of the technology, as has been demonstrated by others ((22), Tombo (bioRxiv doi:10.1101/094672), mCaller (bioRxiv doi:10.1101/127100). Secondly, improvements can be made in reaching true nucleotide-resolution methylation calls. Where multiple CpG sites occur within less than twice the *k*-mer length (here 6) all these sites are considered to have the same methylation state. Again, this is not a limitation of the technology, as more complete training data will allow resolution of mixed methylation states. Finally, we see opportunities for improvement in the analysis of nanopore methylation data. Instead of binary calls from bisulfite sequencing, the output of nanopore sequencing is a likelihood ratio that the site is methylated vs. unmethylated. Currently there is no method for the detection of differential methylation that accepts these continuous values as input. Additionally, the DMR detection algorithm that we used was designed originally for bisulfite data, and we expect that algorithms designed specifically to incorporate long reads and probabilistic methylation assignment would achieve greater levels of accuracy.

## Conclusions

We demonstrate that long-read sequencing using nanopore technology can efficiently generate haplotyped mammalian methylomes. With no additional sample preparation than that routinely used for basic sequencing and with only a mean coverage of about 10X, we identify differential allelic methylation throughout the genome. Combined with expression data, this improves the resolution of imprinting analyses. Our approach is widely applicable to other systems, for instance with more complex genetics, or to phase cancer mutations with methylation state, and to determine the effects of structural variation on methylation.

## Methods

### Animal strains and husbandry

All mice were maintained and treated in accordance with Walter and Eliza Hall Institute Animal Ethics Committee approved protocols under approval number WEHI AEC 2014.026. *Mus musculus castaneus* mice were obtained from Jackson Labs. Note that due to prior inter-crossing for transgene transmission the female *Mus musculus domesticus* C57BL/6 mice that served as dams for our study comprise 12.5% FVB/NJ genome, however for simplicity we will refer to this mouse as B6. Wild-type B6 were reciprocally mated to wild type *Mus musculus castaneus* (Cast).

### DNA and RNA extraction

Pregnant females were sacrificed at E14.5 by CO2 asphyxiation and the embryonic portion of each placenta was dissected from the maternal portion in PBS, as we have done previously (47). Samples were snap frozen in buffer RLT plus (Qiagen) and DNA and RNA were later extracted from the same sample using the AllPrep DNA/RNA Mini Kit (Qiagen), according to manufacturer’s instructions. Samples were sexed by PCR using primers for *Otc* (X-linked gene) and *Zfy* (Y-linked gene) as previously described (48), and male samples were selected for further analysis.

### Illumina sequencing

RRBS libraries were made from 100 ng of DNA purified from the embryonic layer of a male B6 × Cast E14.5 placenta using the Ovation RRBS MethylSeq System (NuGEN), according to the manufacturer’s recommendations, which include use of the Qiagen Epitect kit for bisulfite conversion. The resultant library was sequenced on a HiSeq 2500 (Illumina) using 100 bp paired end reads and analysed as previously described (48, 49).

RNA-seq libraries were prepared from 1 *μ*g of RNA from four B6 × Cast and four Cast ×B6 samples, including the same sample as the RRBS library, using the TruSeq RNA sample preparation kit (Illumina). 75-bp paired-end sequencing was performed on a NextSeq 500 (Illumina). Reads were trimmed with *Trim Galore* v0.4.2 and mapped with *HISAT2* v2.0.5 (50) with option `--no-softclip` to the GRCm38 (mm10) mouse genome with N-masked *castaneus* SNPs. Mapped reads were haplotyped with *SNPsplit* v0.3.2 (31), and gene counts obtained by running *featureCounts* (51) on the GRCm38_v90 Ensembl annotation. Differential analysis was performed with *edgeR* (52, 53) using quasi-likelihood fits (54) and controlling the false discovery rate (FDR) at 10% (55). Interactive plots were produced with Glimma (56).

### Nanopore sequencing

The B6 × Cast F1 sample was sequenced on three MinION flow cells with the 1D Sequencing Genomic Ligation (LSK108) protocol from ONT with minor adjustments: 4 *μ*g of starting material were used for each library preparation, and for two libraries DNA was sheared to 10 kb with a Covaris G-Tube, whereas no shearing was done for the third library (resulting in longer read lengths). Reads were basecalled with Albacore 1.2.2. The Cast × B6 F1 was sequenced on one PromethION flow c ell w ith the 1D Sequencing Genomic Ligation (LSK109) protocol without shearing, and basecalled with Albacore 2.2.7. Nanopore reads were aligned to the same SNP-masked genome as before, using *BWA-MEM* (arXiv:1303.3997).

### Haplotyping

Haplotyping is achieved through the identification of SNPs that are unique to one or the other allele. Examining only the SNPs identified as passing all filters in Keane *et al.* (57), we combine two distinct methods to confidently haplotype each read.

#### Basecall Haplotypin

Where a read is aligned to a SNP position *i* on the reference genome, we assign a score *S*_*i*_ if the aligned base agrees with the reference haplotype, or 1 − *S*_*i*_ if the aligned base agrees with the alternate haplotype, where the score depends on the basecalling quality score *q_i_* of the base in question as

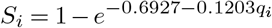

where the coefficients of the above relationship were determined empirically on successfully haplotyped reads. Bases which match neither haplotype, or which exhibit a deletion at the SNP location are excluded from the analysis. Finally, the read is assigned an aggregate haplotype value *h* ∈[0, 1] across the *n* informative SNP calls as follows:

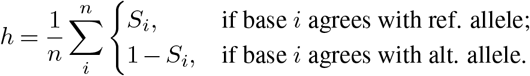

#### Signal-level haplotyping

For signal-level haplotyping, we use the HMM of Simpson *et al.*, implemented in *nanopolish phase-reads* (30). Briefly, the raw signal corresponding to the section of the read aligned to the reference at the SNP position is realigned using a Hidden Markov Model, and the likelihood of the sequence of 6-mers in this vicinity is maximised by choosing the more likely of the two possible alleles. Each read is then assigned scores according to the same rule as in *Basecall Haplotyping*, where *nanopolish* quality scores are offset by −35 in order to exhibit a similar relationship to basecall quality scores.

#### Combining Haplotype Calls

For each read with *n*_*base*_ and *n*_*signal*_ associated SNP calls and associated haplotype values *h*_*base*_ and *h*_*signal*_, we define the haplotype calls

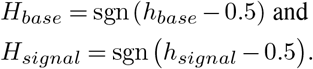

The calls are then combined according to the following rules, applied in order:

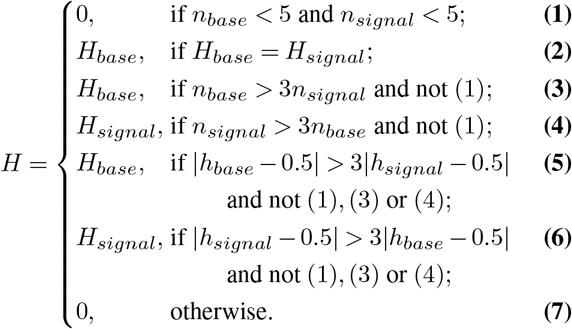

where *H* = 1 represents a read assigned to the reference haplotype, *H* = −1 represents a read assigned to the alternate haplotype, and *H* = 0 represents an unassigned read. This process is shown graphically as a flowchart in Fig. S3.

#### Resolution of maternal recombination

Owing to the cross of an FVB-strain into the maternal line in the grand-parental generation, it is necessary to resolve which section of the maternal genome was contributed by recombination from the FVB chromosome. We run the above haplotyping procedure with three possible outcomes, rather than two: mm10, FVB, and CAST, with variants called by Keane *et al.* (57). The proportion of maternal (non-CAST) reads within any 100Kb region was fitted to are cursive partition tree, which splits continuous data into a stepwise function, here representing the proportion of a contiguous section of chromosome haplotyped to FVB (Fig. S5). Fitting was performed using the R package *rpart* with parameters `minsplit=5` and `cp=0.1` (58). SNPs in sections of the chromosome with mean proportion of FVB greater than 50% were replaced with the FVB allele for further analysis.

### Methylation calling

We determined the methylation status of each CpG site on each read using *nanopolish call-methylation* (21). Briefly, *nanopolish* uses a 5-base alphabet, with 5-methylcytosine represented as M, to build a Gaussian mixture model representing every possible 6-mer with both methylated and unmethylated cytosine in a CpG context, excluding those 6-mers which contain both the methylated and unmethylated base. We ran *nanopolish* separately on reads haplotyped to the maternal and paternal chromosome, using a SNP-masked version of each chromosome to decrease bias in reads with expected deviations from the mm10 reference. *Nanopolish* then assigns each section or “event” of nanopore current to a base on the reference genome and calculates the likelihood of each 6-mer containing the CpG site being either methylated (*L*_*M*_ (*d*_*ij*_)) or unmethylated (*L*_*C*_ (*d*_*ij*_)) given the data *d*_*ij*_ for a call group *i* covered by a read *j*. Groups of consecutive CpG sites in which the distance between any two adjacent sites is less than 11 bases (therefore having overlap between 6-mers containing the cytosines in question) are chained into *CpG call groups*. All sites within the one CpG call group are assumed to have the same methylation status, such that each 6-mer is only considered once. We convert these likelihoods to probabilities as follows:

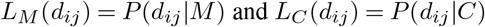

By Bayes’ Law,

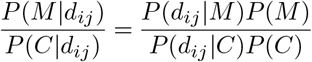

and since *M* and *C* are mutually exclusive and jointly exhaustive,

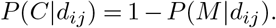

Then, defining the prior probability of methylation as, *P*(*M*) = *p*_0_,

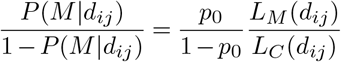

and rearranging for *P* (*M |d_ij_*),

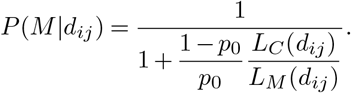

Noting results from Decato *et al.* (28) showing methylation levels ranging from 0.433 to 0.538 for mouse placental tissue we set *p*_0_ = 0.5, so finally we define the single-read, single-site probability of methylation as

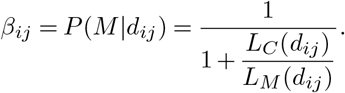

#### Comparison with RRBS methylation calls

Individual methylation calls on a single CpG call group are aggregated over the set of reads covering each group in order to compare with aggregate values provided by bisulfite sequencing. That is, for each CpG call group *i* covered by *n* reads, we define the call group average

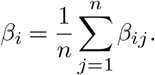

In order to compare methylation calls between nanopore and reduced representation bisulfite sequencing, we must split CpG call groups defined by *nanopolish* as CpG sites separated by less than 11 base pairs into individual sites, including GpC sites on the reverse strand, with each site retaining the same *β* value as the original call group. Only those CpG sites for which both RRBS and nanopore data exist are considered.

### Identification of differentially methylated regions

Following methylation detection and haplotype assignment of each read, it is possible to assign each call of methylation on the genome to one of the two haplotypes. The aggregated *β* methylation values for each CpG group are tested for DMRs using DSS (37). Briefly, DSS tests for differentially methylation at a single CpG-sites, using a Wald test on the coefficients of a beta-binomial regression of count data with an ‘arcsine’ link function. Then, using a default p-value threshold of 10^−5^, DSS aggregates differentially methylated sites into DMRs based on a maximum separation between sites and a minimum density and number of sites in each DMR. To detect parent-of-origin DMRs, we perform DSS with the comparison B6 and Cast vs. Cast and B6; to detect strain-specific D MRs, we perform DSS a second time with the comparison B6♀ and B6♂ vs. Cast♂ and Cast♀.

### Visualisation of haplotyped methylation

Owing to the noisy nature of nanopore methylation calls, we use a loess smoothing curve to visually represent the methylation of a single nanopore read (59). Here the smoothing parameter *α* is determined by

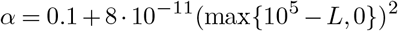

where *L* is read length. This relationship was determined empirically to have minimal impact on visualisation while minimising computation time.

## Data availability

All sequencing data are available at ENA under study accession ERP109201. Processed data can be explored via a Genome browser (60, 61), interactive plots (56) and a summary page available from bioinf.wehi.edu.au/haplotyped_methylome. All analysis scripts are available on GitHub at github.com/scottgigante/haplotyped-methylome.

## List of abbreviations

AUROC: Area Under Receiver Operating Characteristic curve
CpG: 5’-C-phosphate-G-3’
CGI: CpG island
DMR: differentially methylated region
DML: differentially methylated locus
F1: first filial generation
FDR: false discovery rate
ICR: imprinting control region
HMM: Hidden Markov Model
ONT: Oxford Nanopore Technologies
PB: Pacific Biosciences
RRBS: reduced representation bisulfite sequencing
SNP: single nucleotide polymorphism
TSS: transcriptional start site

## AUTHOR CONTRIBUTIONS

Tissue collection was performed by TB and QG. ONT sequencing was performed by AL (with supervision from MT and LG) and QG. RRBS was performed by AK and TB, and RNA-seq by QG. Alignment, methylation calling and haplotyping of nanopore data was performed by SG. AK analysed the RRBS and QG the RNA-seq. Development of visualisation and DMR testing was performed by SG, CW and TPS. Figures were generated by SG, QG and AL. SG, QG, AK, TPS, MEB and MER interpreted the data. SG, QG, CW, MEB and MER wrote the manuscript. All authors read and approved the final version.

## ACKNOWLEDGEMENTS

We would like to thank Jared Simpson for his help running *Nanopolish* and Stephen Wilcox for the Illumina sequencing. MEB was supported by Bellberry-Viertel Senior Medical Research Fellowship. MER was supported by an Australian National Health and Medical Research Council (NHMRC) fellowship (GNT1104924). This work was made possible through NHMRC Project grants GNT1098290 and GNT1140976 (to MEB and MER), Victorian State Government Operational Infrastructure Support and NHMRC Research Institute Infrastructure Support Scheme.

## Additional Files

**Additional file 1 — List of detected imprinted genes.** Tab-separated values containing names, coordinates, log2 fold-change and adjusted p-value for the 135 genes with parent-biased expression in E14.5 embryonic placenta.

**Additional file 2 — List of detected DMRs.** Comma-separated values containing coordinates, differential methylation, type (imprinted or strain-specific), and nearby gene for all 933 *de novo* DMRs.

**Fig. S1.**
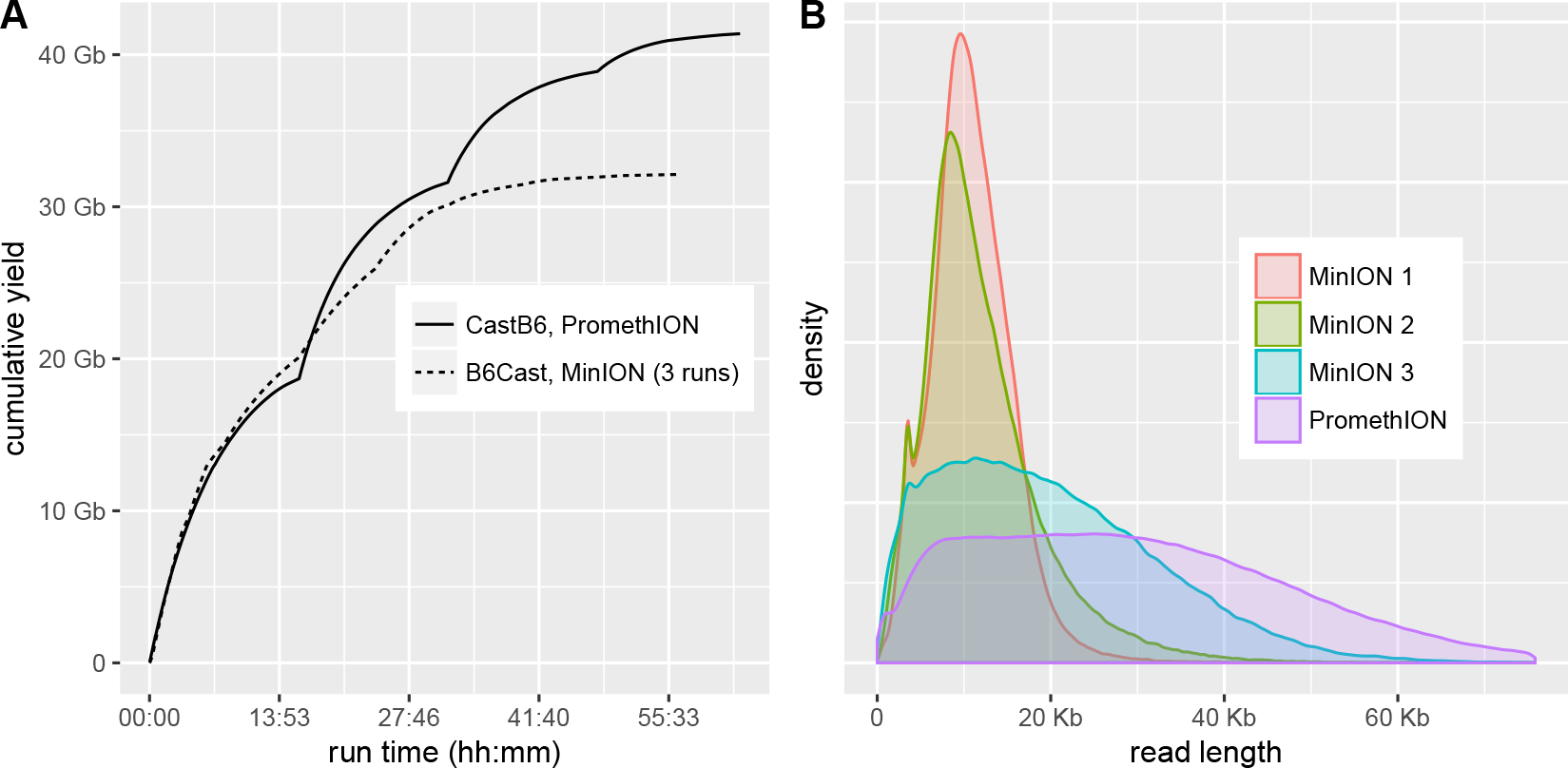
Nanopore sequencing yield and read length. The B6 × Cast (B6Cast) F1 sample was sequenced on three MinION flowcells, and the Cast × B6 (CastB6) F1 on one PromethION flowcell. **A.** Yield of sequencing runs over time (the three MinION runs are merged). **B.** Read length distribution in each sequencing run.

**Fig. S2.**
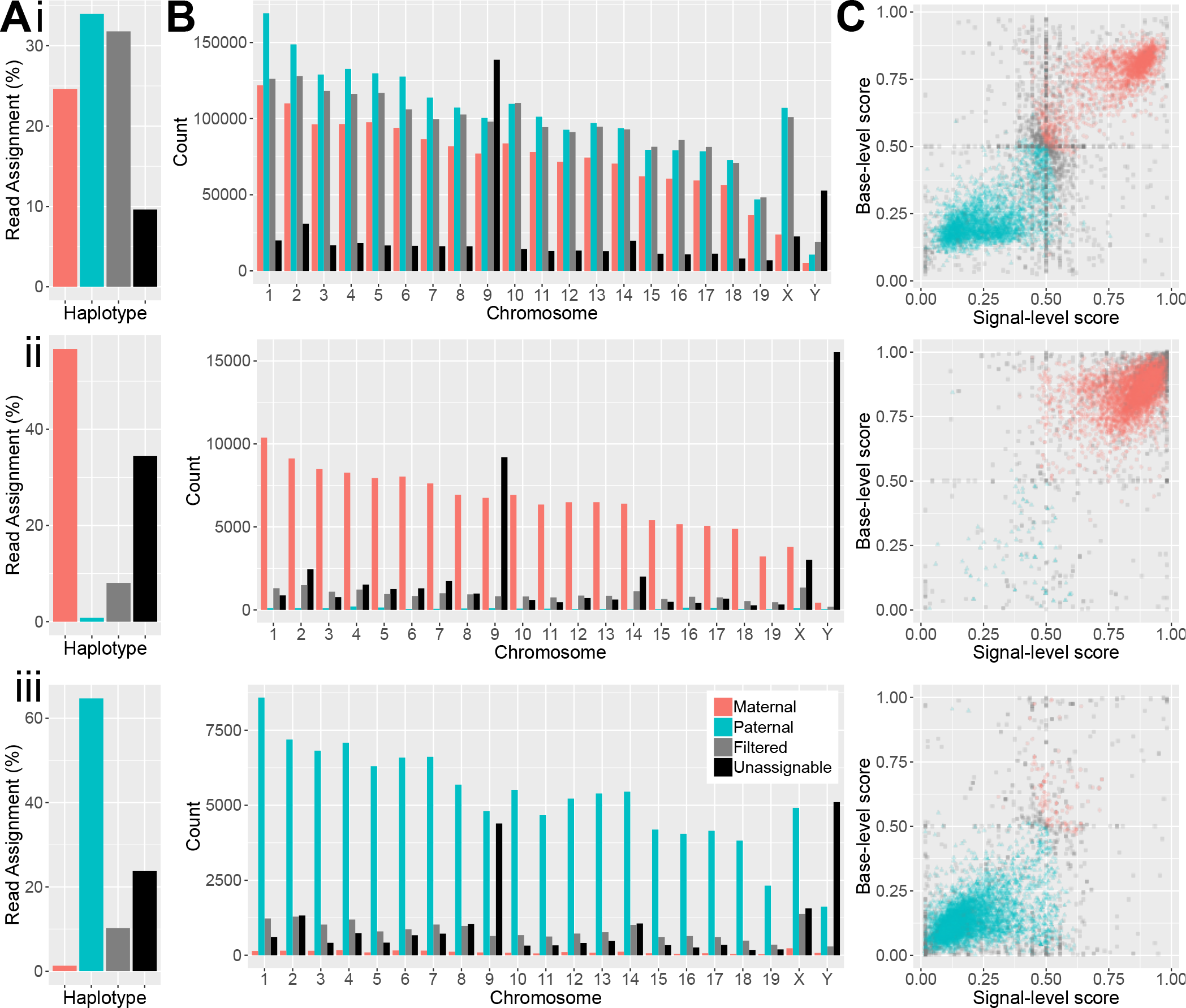
Haplotyping of the Cast × B6, B6 and Cast samples. **A.** Percentages of mapped reads from nanopore sequencing that were assigned to the B6 genome (maternal), Cast genome (paternal), or that could not be haplotyped (filtered) for the Cast × B6 F1 sample (i), *B6* F0 sample (ii) and Cast F0 sample (iii). **B.** Percentages of mapped reads from nanopore sequencing of each sample that were assigned to each haplotype on each chromosome. **C.** Scatter plot of haplotype scores for nanopore reads according to signal (x-axis) and basecall (y-axis) methods. Only 10,000 randomly selected reads are shown for ease of visualisation.

**Fig. S3.**
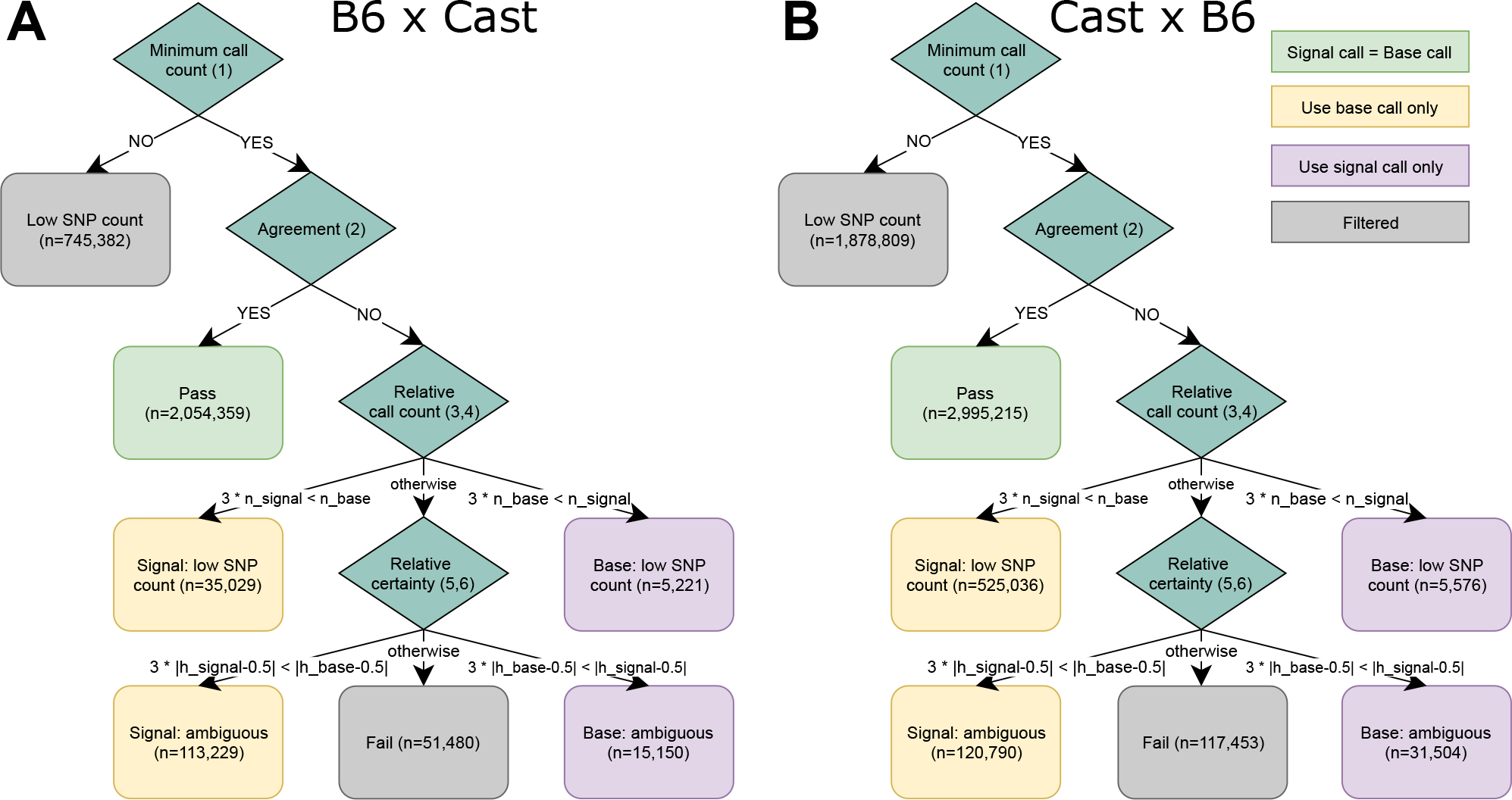
Haplotyping flow chart. **A.** Haplotyping flow chart for B6 × Cast forward cross MinION data. **B.** Haplotyping flow chart for Cast × B6 reverse cross PromethION data.

**Fig. S4.**
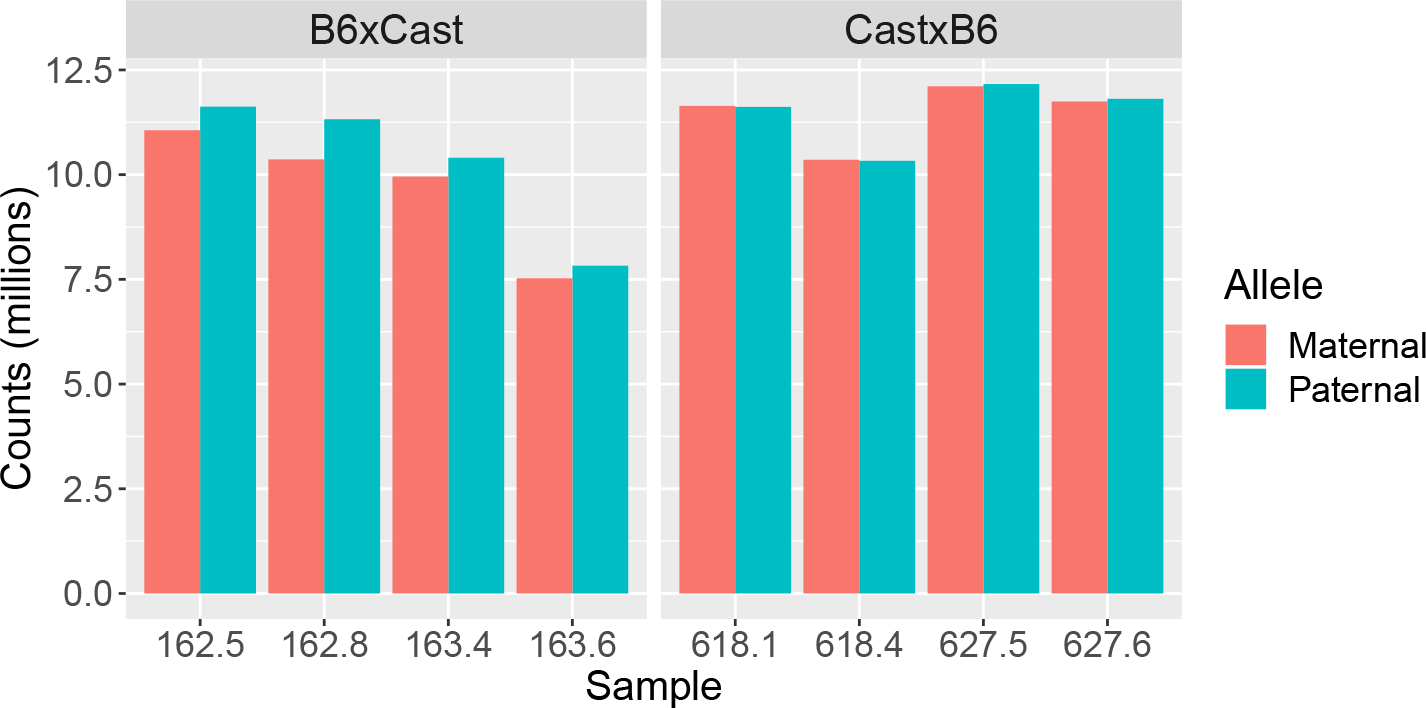
RNA-seq library sizes by allele. Consistent paternal and maternal counts indicate that there was no contamination by maternal tissue. The slightly lower maternal counts in the B6Cast samples are due to the FVB regions in the maternal strain (see Methods).

**Fig. S5.**
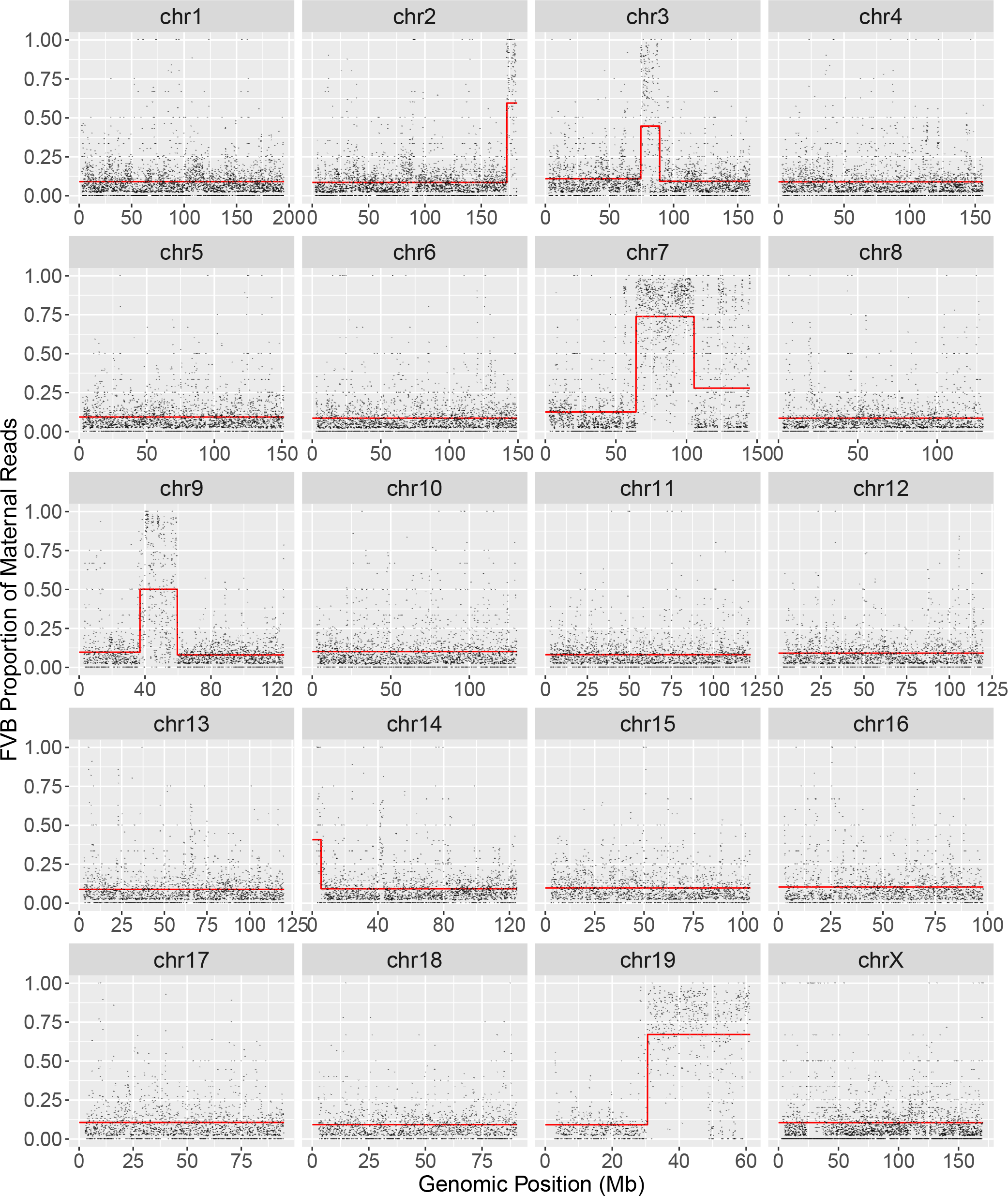
Recursive partition tree resolution of FVB genotype. Proportion of maternal reads from the B6×Cast sample assigned to the FVB genotype by genomic location, where the remainder of the maternal reads are assigned to the C57BL/6 reference genotype. Sections of the genome where the regression tree gave a value higher than 0.4 (shown in red) were assigned to the FVB genotype. The Y chromosome is excluded.

## Bibliography

1. Anne C. Ferguson-Smith, Hiroyuki Sasaki, Bruce M. Cattanach, and M. Azim Surani. Parental-origin-specific epigenetic modification of the mouse h19 gene. Nature, 362 (6422):751–755, 1993. ISSN 0028-0836. doi: 10.1038/362751a0.

2. R Stöger, P Kubicka, C G Liu, T Kafri, A Razin, H Cedar, and D P Barlow. Maternal-specific methylation of the imprinted mouse Igf2r locus identifies the expressed locus as carrying the imprinting signal. Cell, 73(1):61–71, 1993. ISSN 0092-8674. doi: 10.1016/0092-8674(93)90160-R.

3. M S Bartolomei, A L Webber, M E Brunkow, and S M Tilghman. Epigenetic mechanisms underlying the imprinting of the mouse H19 gene. Genes Dev, 7:1663–1673, 1993. ISSN 0890-9369. doi: 10.1101/gad.7.9.1663.

4. En Li, Caroline Beard, and Rudolf Jaenisch. Role for DNA methylation in genomic imprinting. Nature, 366(6453):362–365, 1993. ISSN 00280836. doi: 10.1038/366362a0.

5. Anne C Ferguson-Smith. Genomic imprinting: the emergence of an epigenetic paradigm. Nat Rev Genet, 12(8):565–575, 2011.

6. D. Haig. Coadaptation and conflict, misconception and muddle, in the evolution of genomic imprinting. Heredity, 113(2):96–103, 2014. ISSN 0018-067X. doi: 10.1038/ hdy.2013.97.

7. Jennifer M. Frost and Gudrun E. Moore. The importance of imprinting in the human placenta. PLoS Genet, 6(7):1–9, 2010. ISSN 15537390. doi: 10.1371/journal.pgen.1001015.

8. Denise P Barlow and Marisa S Bartolomei. Genomic Imprinting in Mammals. Cold Spring Harb Perspect Biol, 6(2):a018382–a018382, 2014. ISSN 1943-0264. doi: 10. 1101/cshperspect.a018382.

9. Julio D. Perez, Nimrod D. Rubinstein, and Catherine Dulac. New Perspectives on Genomic Imprinting, an Essential and Multifaceted Mode of Epigenetic Control in the Developing and Adult Brain. Annu Rev Neurosci, 39(1):347–384, 2016. ISSN 0147-006X. doi: 10.1146/annurev-neuro-061010-113708.

10. Xu Wang, Paul D. Soloway, and Andrew G. Clark. A Survey for Novel Imprinted Genes in the Mouse Placenta by mRNA-seq. Genetics, 189(1):109–122, 2011. ISSN 0016-6731. doi: 10.1534/genetics.111.130088.

11. Hiroaki Okae, Hitoshi Hiura, Yuichiro Nishida, Ryo Funayama, Satoshi Tanaka, Hatsune Chiba, Nobuo Yaegashi, Keiko Nakayama, Hiroyuki Sasaki, and Takahiro Arima. Re-investigation and RNA sequencing-based identification of genes with placenta-specific imprinted expression. Hum Mol Genet, 21(3):548–558, 2012. ISSN 1460-2083. doi: 10.1093/hmg/ddr488.

12. Elizabeth H. Finn, Cheryl L. Smith, Jesse Rodriguez, Arend Sidow, and Julie C. Baker. Maternal bias and escape from X chromosome imprinting in the midgestation mouse placenta. Dev Biol, 390(1):80–92, 2014. ISSN 1095564X. doi: 10.1016/j.ydbio.2014.02.020.

13. J Mauro Calabrese, Joshua Starmer, Megan D Schertzer, Della Yee, and Terry Magnuson. A Survey of Imprinted Gene Expression in Mouse Trophoblast Stem Cells. G3 (Bethesda), 5(5):751–759, 2015. ISSN 2160-1836. doi: 10.1534/g3.114.016238.

14. Azusa Inoue, Lan Jiang, Falong Lu, Tsukasa Suzuki, and Yi Zhang. Maternal H3K27me3 controls DNA methylation-independent imprinting. Nature, 547(7664): 419–424, 2017. ISSN 0028-0836. doi: 10.1038/nature23262.

15. Daniel Andergassen, Christoph P Dotter, Daniel Wenzel, Verena Sigl, Philipp C Bammer, Markus Muckenhuber, Daniela Mayer, Tomasz M Kulinski, Hans-Christian Theussl, Josef M Penninger, Christoph Bock, Denise P Barlow, Florian M Pauler, and Quanah J Hudson. Mapping the mouse Allelome reveals tissue-specific regulation of allelic expression. eLife, 6:1–29, 2017. ISSN 2050-084X. doi: 10.7554/eLife.25125.

16. Melanie A. Eckersley-Maslin, Celia Alda-Catalinas, and Wolf Reik. Dynamics of the epigenetic landscape during the maternal-to-zygotic transition. Nat Rev Mol Cell Biol, 19(7):436–450, 2018. ISSN 1471-0072. doi: 10.1038/s41580-018-0008-z.

17. Rachel Duffié, Sophie Ajjan, Maxim V. Greenberg, Natasha Zamudio, Martin del Arenal Escamilla, Julian Iranzo, Ikuhiro Okamoto, Sandrine Barbaux, Patricia Fauque, and Déborah Bourc’his. The Gpr1/Zdbf2 locus provides new paradigms for transient and dynamic genomic imprinting in mammals. Genes Dev, 28(5):463–478, 2014. ISSN 08909369. doi: 10.1101/gad.232058.113.

18. Elmar Schilling, Carol El Chartouni, and Michael Rehli. Allele-specific DNA methylation in mouse strains is mainly determined by cis-acting sequences. Genome Res, 19(11): 2028–2035, 2009. doi: 10.1101/gr.095562.109.

19. M. P. Hitchins, R. W. Rapkins, C. T. Kwok, S. Srivastava, J. J. Wong, L. M. Khachigian, P. Polly, J. Goldblatt, and R. L. Ward. Dominantly inherited constitutional epigenetic silencing of MLH1 in a cancer-affected family is linked to a single nucleotide variant within the 5’UTR. Cancer Cell, 20(2):200–213, 2011. doi: 10.1016/j.ccr.2011.07.003.

20. Benjamin A. Flusberg, Dale R. Webster, Jessica H. Lee, Kevin J. Travers, Eric C. Olivares, Tyson A. Clark, Jonas Korlach, and Stephen W. Turner. Direct detection of DNA methylation during single-molecule, real-time sequencing. Nat Methods, 7(6): 461–465, 2010. ISSN 1548-7091. doi: 10.1038/nmeth.1459.

21. Jared T. Simpson, Rachael E. Workman, P.C. Zuarte, Matei David, L.J. Dursi, and Winston Timp. Detecting DNA methylation using nanopore sequencing. Nat Methods, 14(4):407–410, 2017. doi: 10.1038/nmeth.4184.

22. Arthur C Rand, Miten Jain, Jordan M Eizenga, Audrey Musselman-Brown, Hugh E Olsen, Mark Akeson, and Benedict Paten. Mapping DNA methylation with high-throughput nanopore sequencing. Nat Methods, 14(4):411–413, 2017. ISSN 1548-7091. doi: 10.1038/nmeth.4189.

23. Pacific Biosciences. Detecting DNA base modifications using single molecule, real-time sequencing. Pacific Biosciences, 2012.

24. Y. Yang, R. Sebra, B. S. Pullman, W. Qiao, I. Peter, R. J. Desnick, C. R. Geyer, J. F. DeCoteau, and S. A. Scott. Quantitative and multiplexed dna methylation analysis using long-read single-molecule real-time bisulfite sequencing (SMRT-BS). BMC Genomics, 16:350, 2015.

25. Anthony Rhoads and Kin Fai Au. Pacbio sequencing and its applications. Genomics Proteomics Bioinformatics, 13(5):278–289, 2015.

26. Miten Jain, Sergey Koren, Karen H Miga, Josh Quick, Arthur C Rand, Thomas A Sasani, John R Tyson, Andrew D Beggs, Alexander T Dilthey, Ian T Fiddes, et al. Nanopore sequencing and assembly of a human genome with ultra-long reads. Nature biotechnology, 36(4):338, 2018.

27. R. A. Irizarry, C. Ladd-Acosta, B. Wen, Z. Wu, C. Montano, P. Onyango, H. Cui, K. Gabo, M. Rongione, M. Webster, H. Ji, J. Potash, S. Sabunciyan, and A. P. Feinberg. The human colon cancer methylome shows similar hypo- and hypermethylation at conserved tissue-specific CpG island shores. Nat Genet, 41(2):178–186, 2009. doi: 10.1038/ng.298.

28. Benjamin E Decato, Jorge Lopez-Tello, Amanda N Sferruzzi-Perri, Andrew D Smith, and Matthew D Dean. DNA methylation divergence and tissue specialization in the developing mouse placenta. Mol Biol Evol, 34(7):1702–1712, 2017. doi: 10.1093/ molbev/msx112.

29. Denise L Walker, Aditya Vijay Bhagwate, Saurabh Baheti, Regenia L Smalley, Christopher A Hilker, Zhifu Sun, and Julie M Cunningham. Dna methylation profiling: comparison of genome-wide sequencing methods and the infinium human methylation 450 bead chip. Epigenomics, 7(8):1287–1302, 2015.

30. Nicholas J Loman, Joshua Quick, and Jared T Simpson. A complete bacterial genome assembled de novo using only nanopore sequencing data. Nat Methods, 12(8):733–735, 2015.

31. Felix Krueger and Simon R Andrews. SNPsplit: Allele-specific splitting of alignments between genomes with known SNP genotypes. F1000Res, 5:1479, 2016. ISSN 2046-1402. doi: 10.12688/f1000research.9037.1.

32. Jerome Friedman, Trevor Hastie, and Rob Tibshirani. glmnet: Lasso and elastic-net regularized generalized linear models. R package version, 1(4), 2009.

33. Ian M Morison, Joshua P Ramsay, and Hamish G Spencer. A census of mammalian imprinting. Trends Genet, 21(8):457–465, 2005. doi: 10.1016/j.tig.2005.06.008.

34. RL Jirtle. World Wide Web Site - Geneimprint Imprinted genes database, 2001.

35. Reiner Schulz, Kathryn Woodfine, Trevelyan R. Menheniott, Deborah Bourc’his, Timothy Bestor, and Rebecca J. Oakey. WAMIDEX: A web atlas of murine genomic imprinting and differential expression. Epigenetics, 3(2):89–96, 2008. ISSN 1559-2294. doi: 10.4161/epi.3.2.5900.

36. CM Williamson, A Blake, S Thomas, CV Beechey, J Hancock, BM Cattanach, and J Peters. World Wide Web Site - Mouse Imprinting Data and References. Oxfordshire: MRC Harwell, 2013.

37. Yongseok Park and Hao Wu. Differential methylation analysis for bs-seq data under general experimental design. Bioinformatics, 32(10):1446–53, 2016. doi: 10.1093/ bioinformatics/btw026.

38. Hisato Kobayashi, Takayuki Sakurai, Misaki Imai, Nozomi Takahashi, Atsushi Fukuda, Obata Yayoi, Shun Sato, Kazuhiko Nakabayashi, Kenichiro Hata, Yusuke Sotomaru, Yutaka Suzuki, and Tomohiro Kono. Contribution of Intragenic DNA Methylation in Mouse Gametic DNA Methylomes to Establish Oocyte-Specific Heritable Marks. PLoS Genet, 8(1):e1002440, 2012. ISSN 1553-7404. doi: 10.1371/journal.pgen.1002440.

39. Sébastien A. Smallwood, Shin Ichi Tomizawa, Felix Krueger, Nico Ruf, Natasha Carli, Anne Segonds-Pichon, Shun Sato, Kenichiro Hata, Simon R. Andrews, and Gavin Kelsey. Dynamic CpG island methylation landscape in oocytes and preimplantation embryos. Nat Genet, 43(8):811–814, 2011. ISSN 10614036. doi: 10.1038/ng.864.

40. Zachary D. Smith,, Michelle M. Chan, Tarjei S. Mikkelsen, Hongcang Gu, Andreas Gnirke, Aviv Regev, and Alexander Meissner. A unique regulatory phase of DNA methylation in the early mammalian embryo. Nature, 484(7394):339–344, 2012. ISSN 0028-0836. doi: 10.1038/nature10960.

41. T. Moore, M. Constancia, M. Zubair, B. Bailleul, R. Feil, H. Sasaki, and W. Reik. Multiple imprinted sense and antisense transcripts, differential methylation and tandem repeats in a putative imprinting control region upstream of mouse Igf2. Proc Natl Acad Sci U S A, 94(23):12509–12514, 1997. ISSN 0027-8424. doi: 10.1073/pnas.94.23.12509.

42. Hiroaki Okae, Shogo Matoba, Takeshi Nagashima, Eiji Mizutani, Kimiko Inoue, Narumi Ogonuki, Hatsune Chiba, Ryo Funayama, Satoshi Tanaka, Nobuo Yaegashi, Keiko Nakayama, Hiroyuki Sasaki, Atsuo Ogura, and Takahiro Arima. RNA sequencing-based identification of aberrant imprinting in cloned mice. Hum Mol Genet, 23(4): 992–1001, 2014. ISSN 1460-2083. doi: 10.1093/hmg/ddt495.

43. Simao Teixeira da Rocha, Carol A. Edwards, Mitsuteru Ito, Tsutomu Ogata, and Anne C. Ferguson-Smith. Genomic imprinting at the mammalian Dlk1-Dio3 domain. Trends Genet, 24(6):306–316, 2008. ISSN 01689525. doi: 10.1016/j.tig.2008.03.011.

44. Rachel J Smith, Wendy Dean, Galia Konfortova, and Gavin Kelsey. Identification of Novel Imprinted Genes in a Genome-Wide Screen for Maternal Methylation. Genome Res, 13(4):558–569, 2003. ISSN 10889051. doi: 10.1101/gr.781503.

45. Franck Court, Chiharu Tayama, Valeria Romanelli, A. Martin-Trujillo, Isabel Iglesias-Platas, Kohji Okamura, Naoko Sugahara, C. Simon, Harry Moore, Julie V Harness, Hans Keirstead, J. V. Sanchez-Mut, Elsuke Kaneki, Pablo Lapunzina, Hidenobu Soejima, Norio Wake, Manel Esteller, Tsutomu Ogata, Kenichiro Hata, Kazuhiko Nakabayashi, and David Monk. Genome-wide parent-of-origin DNA methylation analysis reveals the intricacies of human imprinting and suggests a germline methylation-independent mechanism of establishment. Genome Res, 24(4):554–569, 2014. ISSN 1088-9051. doi: 10.1101/gr.164913.113.

46. Noriaki Takemoto, Takuji Yoshimura, Satsuki Miyazaki, Fumi Tashiro, and Junichi Miyazaki. Gtsf1l and Gtsf2 Are specifically expressed in gonocytes and spermatids but are not essential for spermatogenesis. PLoS ONE, 11(3):1–12, 2016. ISSN 19326203. doi: 10.1371/journal.pone.0150390.

47. Arne W Mould, Zhenyi Pang, Miha Pakusch, Ian D Tonks, Mitchell Stark, Dianne Carrie, Pamela Mukhopadhyay, Annica Seidel, Jonathan J Ellis, Janine Deakin, Matthew J Wakefield, Lutz Krause, Marnie E Blewitt, and Graham F Kay. Smchd1 regulates a subset of autosomal genes subject to monoallelic expression in addition to being critical for X inactivation. Epigenetics Chromatin, 6:19, 2013.

48. Andrew Keniry, Linden J Gearing, Natasha Jansz, Joy Liu, Aliaksei Z Holik, Peter F Hickey, Sarah A Kinkel, Darcy L Moore, Kelsey Breslin, Kelan Chen, Ruijie Liu, Catherine Phillips, Miha Pakusch, Christine Biben, Julie M Sheridan, Benjamin T Kile, Catherine Carmichael, Matthew E Ritchie, Douglas J Hilton, and Marnie E Blewitt. Setdb1-mediated H3K9 methylation is enriched on the inactive X and plays a role in its epigenetic silencing. Epigenetics Chromatin, 9:16, 2016.

49. Danqing Yin, Matthew E Ritchie, Jafar S Jabbari, Tamara Beck, Marnie E Blewitt, and Andrew Keniry. High concordance between illumina HiSeq2500 and NextSeq500 for reduced representation bisulfite sequencing (RRBS). Genom Data, 10:97–100, 2016. doi: 10.1016/j.gdata.2016.10.002.

50. Daehwan Kim, Ben Langmead, and Steven L Salzberg. HISAT: a fast spliced aligner with low memory requirements. Nat Methods, 12(4):357–60, 2015.

51. Y. Liao, G. K. Smyth, and W. Shi. featureCounts: an efficient general-purpose program for assigning sequence reads to genomic features. Bioinformatics, 30(7):923–30, 2014.

52. M. D. Robinson, D. J. McCarthy, and G. K. Smyth. edgeR: a Bioconductor package for differential expression analysis of digital gene expression data. Bioinformatics, 26: 139–140, 2010.

53. Davis J. McCarthy, Yunshun Chen, and Gordon K. Smyth. Differential expression analysis of multifactor RNA-Seq experiments with respect to biological variation. Nucleic Acids Res, 40(10):4288–4297, 2012. ISSN 03051048. doi: 10.1093/nar/gks042.

54. A. T. Lun, Y. Chen, and G. K. Smyth. It’s DE-licious: A Recipe for Differential Expression Analyses of RNA-seq Experiments Using Quasi-Likelihood Methods in edgeR. Methods Mol Biol, 1418:391–416, 2016. doi: 10.1007/978-1-4939-3578-9$\_$19.

55. Y. Benjamini and Y. Hochberg. Controlling the false discovery rate: a practical and powerful approach to multiple testing. Journal of the Royal Statistical Society: Series B, 57:289–300, 1995.

56. Shian Su, Charity W. Law, Casey Ah-Cann, Marie-Liesse Asselin-Labat, Marnie E. Blewitt, and Matthew E. Ritchie. Glimma: interactive graphics for gene expression analysis. Bioinformatics, 33(13):2050–2052, 2017. ISSN 1367-4803. doi: 10.1093/ bioinformatics/btx094.

57. Thomas M Keane, Leo Goodstadt, Petr Danecek, Michael A White, Kim Wong, Binnaz Yalcin, Andreas Heger, Avigail Agam, Guy Slater, Martin Goodson, et al. Mouse genomic variation and its effect on phenotypes and gene regulation. Nature, 477 (7364):289–94, 2011.

58. Terry Therneau, Beth Atkinson, and Brian Ripley. rpart: Recursive Partitioning and Regression Trees, 2017. R package version 4.1-11.

59. William S Cleveland, Eric Grosse, and William M Shyu. Local regression models. Statistical models in S, 2:309–376, 1992.

60. Robert Buels, Eric Yao, Colin M. Diesh, Richard D. Hayes, Monica Munoz-Torres, Gregg Helt, David M. Goodstein, Christine G. Elsik, Suzanna E. Lewis, Lincoln Stein, and Ian H. Holmes. JBrowse: a dynamic web platform for genome visualization and analysis. Genome Biol, 17(1):66, 2016. ISSN 1474-760X. doi: 10.1186/s13059-016-0924-1;.

61. Brigitte T. Hofmeister and Robert J. Schmitz. Enhanced JBrowse plugins for epigenomics data visualization. BMC Bioinformatics, 19(1):159, 2018. ISSN 1471-2105. doi: 10.1186/s12859-018-2160-z.

